# Invariant relationship unites REM and NonREM sleep

**DOI:** 10.1101/2023.05.08.539925

**Authors:** Irina V. Zhdanova, Vasili Kharchenko

## Abstract

Establishing structural and functional links between two distinct types of sleep, rapid-eye-movement (REM) and non-REM (NREM), that alternate and form several sleep cycles per night, has posed a significant challenge. Here we demonstrate a simple invariant relationship where the product of the duration of NREM sleep episode and intensity of subsequent REM sleep episode remains constant over successive sleep cycles of normal human sleep. This Sleep Cycle Invariant (SCI), previously predicted by a quantitative model of sleep dynamics, supports the structural and functional unity of NREM and REM sleep. The significance of SCI for understanding normal sleep and sleep disorders is highlighted by alterations in REM sleep intensity and NREM sleep episode duration being a hallmark of major depression.

**One-Sentence Summary:** The duration of NREM sleep and intensity of REM sleep have an invariant relationship across normal sleep cycles of one night.

## Main Text

Within the global Sleep-Wake homeostasis (*1*), the principal physiological function of sleep state and its intrinsic dynamics are still a matter of debate (*2, 3*). The issue is further complicated by the existence of two distinct types of sleep: non-rapid-eye-movement sleep (NREMS) and rapid-eye-movement sleep (REMS) (*4*). Besides perceptual and behavioral disengagement from the environment, these two states are strikingly different. Normal sleep dynamics involve orderly alternations of these two types of sleep in episodes of varying duration and intensity.

Sleep starts with NREMS, followed by REMS, together forming one sleep cycle, with adult humans typically experiencing 5-6 cycles per night (Fig. 1A). The cycles are quasi-periodic, with the duration and intensity of NREMS and REMS changing over consecutive cycles in a distinct manner (Fig. 1B). The NREMS is associated with slow brain activity and reduced muscle tone. In REMS, also known as the paradoxical sleep, the wake-like brain activity of fast low amplitude waves and active dream mentation, rapid eye movements, sexual arousal, irregular heart rate and respiration overlaps with further reduction of perception, skeletal muscle paralysis and loss of thermoregulation (*4*). This unique simultaneous presence of wake-like and sleep-like features makes the nature and significance of REMS particularly puzzling.

**Figure 1.**
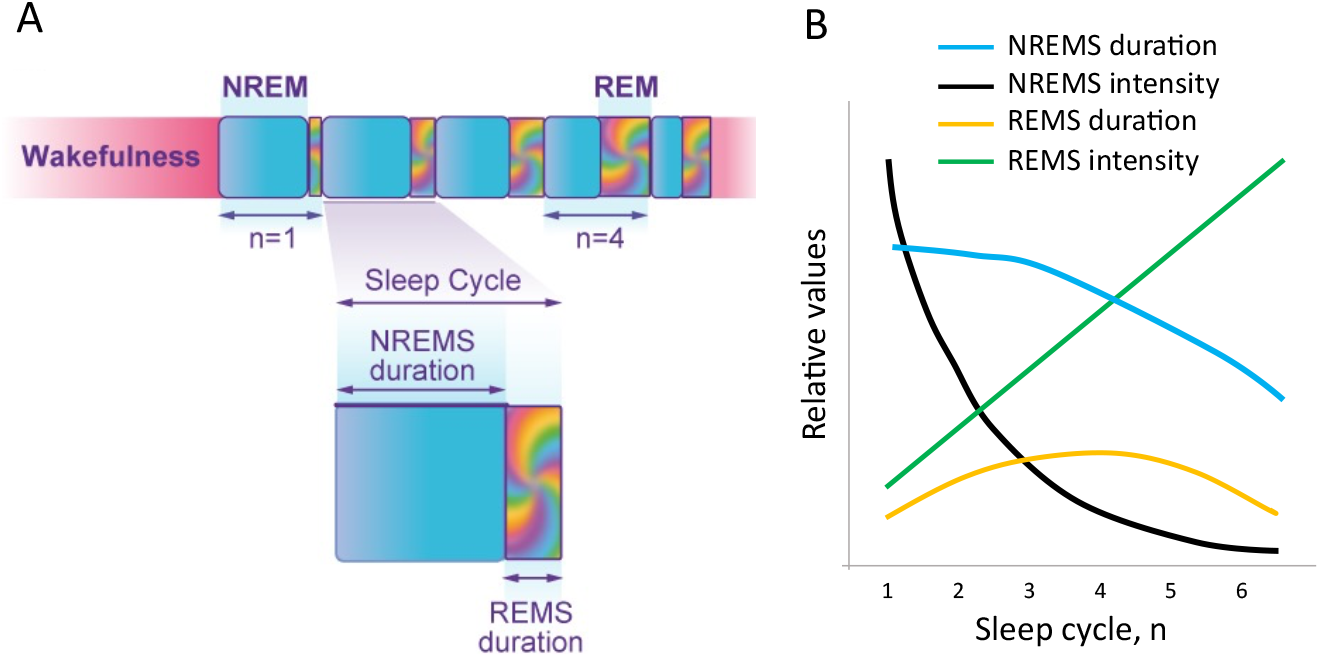
Dynamic structure of human sleep. A. Schematics of consolidated sleep pattern, consisting of consecutive sleep cycles, each including longer NREMS (cyan) and shorter REMS (multicolor) episodes. Five sleep cycles are shown, n. B. Schematics of typical overnight dynamics in the four primary sleep measures over consecutive sleep cycles: NREMS duration (cyan), NREMS intensity (SWS, black), REMS duration (orange) and REMS intensity (REM density, green). From figure 1 in (*9*), with authors’ permission.

Since the discovery of REM sleep was reported in *Science* 70 years ago (*5*), it has been debated whether NREMS and REMS are two essential elements of the same process of sleep homeostasis, dependent on each other, or whether they adaptively coexist through alternation but serve different functions, and are regulated independently (6-8). This debate highlights the need for a unifying conceptual and mathematical model of sleep dynamics that can clarify the relationship between NREMS and REMS. Several model approaches have been applied to address this question, but uncertainties persist (*8*).

Recently, we have proposed a quantitative model of sleep dynamics that suggests a structural and functional unity of NREM and REM sleep (*9*). Our model predicted an invariant relationship between NREMS and REMS, which we named the Sleep Cycle Invariant (SCI). Specifically, the model predicts that the product of NREM sleep duration and REM sleep intensity should remain constant over consecutive sleep cycles. Here we show the experimental evidence of the existence of the SCI.

## Results

### Invariant relationship between the two types of sleep

To determine if the predicted invariant relationship between NREMS and REMS is observed in experimental data, we analyzed the primary sleep measures in young, healthy individuals with high sleep efficiency by assessing NREMS duration and REMS intensity per each sleep cycle. To document REMS intensity, the original quantitative method of counting all the individual rapid eye movements was used to determine the number of movements per minute of REMS episode, i.e., REM density (*10*). The product of NREMS duration and REMS intensity was calculated for each sleep cycle of each individual night to evaluate the SCI.

As expected, in this group, the NREMS duration and REMS intensity showed distinct dynamics over consecutive sleep cycles. However, the product of these two sleep measures (SCI) remained near constant over the course of the night (Fig. 2A,B, P value > 0.92, see Methods), confirming the invariant relationship between NREMS and REMS over consecutive sleep cycles of normal high-efficiency sleep.

**Figure 2.**
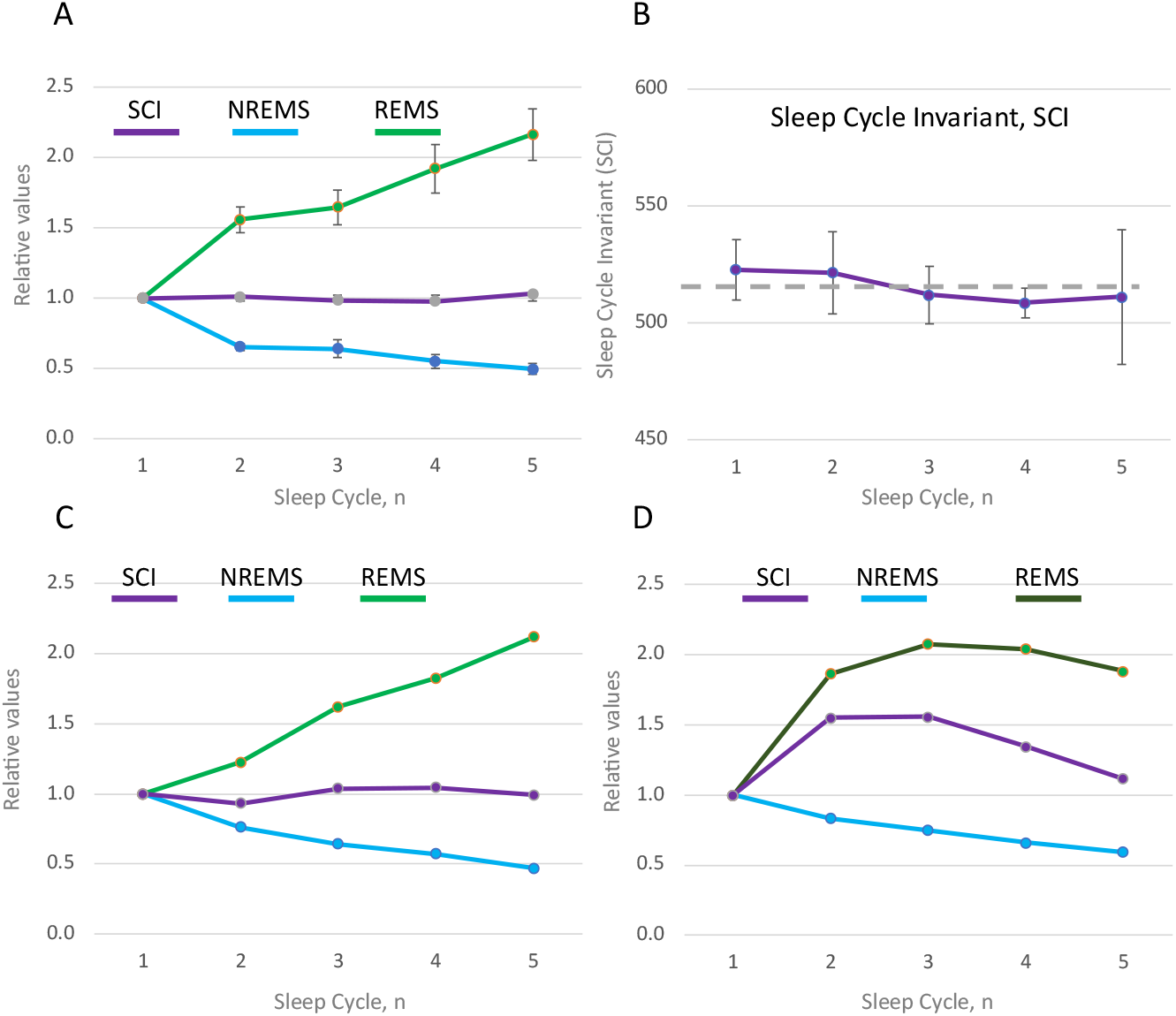
Sleep Cycle Invariant unites NREM and REM sleep. A. The product of NREMS episode duration and REMS episode intensity - the Sleep Cycle Invariant (SCI, purple), NREMS duration (cyan) and REMS intensity based on quantitative evaluation of REM density (light green) over consecutive cycles of regular sleep in a group of 11 young healthy volunteers with sleep efficiency of at least 93%; n=11 for cycles 1-4 and n=5 for cycle 5 (see Methods). Data normalized to the first sleep cycle (=1) for each parameter and presented as group means±SEM per cycle *. B.Variation in SCI over consolidated high-quality sleep. Data are presented as mean ±SEM; dashed line defines the mean value (515.2); the rest as in A. C. Analogous to panel A, the plot shows SCI based on NREMS duration per sleep cycle, as reported by Barbato & Wehr (*15*) (n=308 nights; 11 young healthy volunteers), and REMS intensity per cycle, as reported by Aserinsky (*12*) and evaluated using quantitative method (n=20 nights; 10 young healthy volunteers). D. Loss of linear time- and cycle-dependency of REMS intensity when assessed using semi-quantitative method (dark green) results in obscured NREMS-REMS invariant relationship (purple). REMS intensity was assessed in parallel with NREMS duration, as reported by Barbato et al. (*16*) (n=208 nights; 8 young healthy volunteers). Data normalized to the first sleep cycle (=1) for each parameter and presented as group mean per cycle*. * In A-D, the duration of the first NREMS episode was multiplied by 1.33, as per the model prediction (*9*) of incomplete period of the first cycle (see Methods).

### Revealing the Sleep Cycle Invariant Requires Precise Quantitative Assessment of REMS Intensity

We then aimed to evaluate SCI in previously published data on normal sleep and sleep disorders, but encountered a methodological challenge. Sleep studies typically include three out of the four principal measures of sleep: the duration and intensity of NREM sleep and the duration of REM sleep. However, reports on the intensity of REM sleep, especially based on its quantitative evaluation, are exceptionally rare. This is mainly due to the complexity of the bursts of rapid eye movements (REMs), which makes it difficult to count individual movements both visually and automatically. Instead, many studies employed one of several semi-quantitative assessments, such as assigning a score to a range of REMs or counting number of intervals that contained REMs. These semi-quantitative measures can reveal major pathological changes in REMS intensity but lack sensitivity, especially at high intensity levels (*11*). For this reason, they may obscure normal linear increase in REMS intensity over the sleep period, which was demonstrated through quantitative assessment and manifests independent of the circadian phase (*10-14*). Consequently, we could identify only few studies that reported quantitative data on REMS intensity, and none that reported it over consecutive sleep cycles in parallel with NREMS episode durations.

We then tried a different approach to investigate the SCI. In a previous study, we first predicted and then confirmed that the dynamics of REMS intensity are nearly identical among groups of individuals with normal sleep of comparable habitual duration (*9*). Based on this finding, we aimed to determine whether the SCI could still be observed using sleep data from different sources, provided that the groups studied were similar in age, had normal sleep of typical habitual duration, and REMS intensity was assessed quantitatively by counting each individual eye movement. In this “hybrid” analysis, we used NREMS data from one study (*15*) and REMS intensity data from another (*12*), with both studies being of the highest quality and allowing subjects abundant sleep opportunity. This analysis also revealed the invariant relationship between NREMS and REMS (Fig. 2C). In contrast, when we used the results of semi-quantitative assessment of REMS intensity in combination with NREMS episode duration documented within the same large-scale study of top quality (*16*), the invariant relationship was obscured (Fig. 2D). Together, these results further support the existence of SCI in normal sleep and underscore the critical importance of quantitative assessment of REMS intensity.

## Discussion

The experimental confirmation of the Sleep Cycle Invariant provides compelling evidence that NREMS and REMS are integral parts of a unified process. This finding opens up new avenues for experimentation and conceptual analysis. In physics and other fields, invariant or symmetry relationships are generally indicative of conservation laws and principles governing the behavior, and commonly provide insight into the underlying mechanisms of the system. For instance, in our wave-based model of sleep (*9*), the invariant relationship expressed by SCI arose from a general property of quasi-classical potentials, where the period of wavepacket oscillations is inversely proportional to the gap between energy levels, with the period and energy lost correlating with NREMS duration and REMS intensity, respectively. Other models of sleep dynamics may provide alternative explanations for this phenomenon. Nonetheless, the existence of an invariant relationship between NREMS and REMS strongly suggests that these two types of sleep are not independent but rather regulated by a common mechanism.

The SCI may provide a unique metric that integrates NREMS and REMS to measure sleep quality for each cycle and across the entire sleep period. This presents a significant opportunity to investigate the dynamic mechanisms underlying sleep disturbances, such as the different types of insomnia that affect sleep onset, maintenance, and early morning awakening in distinct ways. Moreover, the SCI may also provide new insights into the pharmacodynamics of both established and novel medications, as they exhibit unique, time-dependent effects on sleep.

The SCI may be particularly relevant in the context of psychiatric and neurological disorders, where sleep disturbances are prevalent and often the first symptom of disease or its relapse (*17-20*). Remarkably, changes in the two measures that form the SCI, the REMS intensity and NREMS duration, are the most prominent in these disorders and found to correlate with disease severity and treatment outcomes (*17-26*). This is especially well-documented for major depression, where these changes are widely accepted as a diagnostic biomarker in patients (*17-22*) and suggested as a vulnerability marker in family members (*23*).

Tendency for preserving the SCI may explain why the changes in these two parameters in pathological conditions are typically coordinated, no matter which direction the shift is (*17-26*). For instance, in affective disorders, an increase in REMS intensity is typically accompanied by a decrease in NREMS episode duration, particularly well documented in the first sleep cycle and often referred to as short latency to REMS (*17-25*). In contrast, patients with Parkinson’s disease exhibit reduced REMS intensity and an increased duration of the first NREMS episode (*26*). Quantitative assessment of REMS intensity over the entire sleep period should determine the extent to which the invariant relationship between NREMS and REMS is preserved or altered in these disorders.

In conclusion, our experimental findings provide empirical evidence of a quantitative invariant relationship between NREM and REM sleep, supporting their intrinsic unity, as predicted by the wave model of sleep dynamics (*9*). The observation of this surprisingly simple connection between the two phenomenologically distinct types of sleep holds significant implications for understanding the overall sleep process and addressing primary sleep disorders, as well as those associated with neurological and psychiatric conditions.

## Acknowledgments

We thank Peter Kharchenko for constructive discussions, valuable advice and statistical analysis, and our entire research team at Boston University School of Medicine for technical assistance in data collection.

## Funding

Chaikin-Wile Foundation (VK)

Pfizer Inc. (IVZ)

Biochron LLC (IVZ)

## Author contributions

Conceptualization: VK, IVZ

Methodology: VK

Investigation: IVZ

Visualization: VK, IVZ

Funding acquisition: VK, IVZ

Project administration: VK, IVZ

Supervision: VK, IVZ

Writing – original draft: VK, IVZ

Writing – review & editing: VK, IVZ

## Data and materials availability

The group sleep data results for NREMS duration and REMS intensity used in the current analysis of SCI are available from the corresponding author on reasonable request.

## Supplementary materials

Materials and Methods

## Materials and Methods

### (a) Experimental Design

The data presented here (Fig. 2A,B) were part of our larger study on the circadian regulation of sleep and hormonal functions (“Multimodal Circadian Rhythm Evaluation” PI: IVZ), which will be reported in full elsewhere. The study was conducted in accordance with the Declaration of Helsinki on Ethical Principles for Medical Research Involving Human Subjects, adopted by the General Assembly of the World Medical Association, and approved by the Boston University Institutional Review Board. All the participants provided written informed consent.

#### Subjects

The subjects whose data were analyzed for the assessment of SCI were 11 of the overall group of 24 young healthy male volunteers (mean ±SEM: 23.5 ± 2.1 years of age, ranging 19-31 years of age) who, along with the rest of the subjects, were selected based on the following self-reported criteria: 7-9 hours of habitual nighttime sleep, small (<1.5h) changes in sleep length on weekends, no sleep complaints, no history of chronic disorders or regular medications, no recent trans-meridian travel, no drug use, no smoking, habitual coffee consumption not exceeding 3 cups a day.

#### Experimental protocol

Over the two weeks prior to the inpatient part of the study, the sleep-wake cycle was documented using activity monitors (Phillips Inc.) and a sleep log. Starting on Friday night, subjects spent 3 consecutive nights in the General Clinical Research Center of Boston University School of Medicine. The time in bed was scheduled individually to correspond to the habitual bedtime and subjects were allowed to stay in bed for 9 consecutive hours. Sleep was recorded using polysomnography (Nihon Kohden PSG system), as per standard techniques, and the sleep stages were visually scored for consecutive 30-s epochs (*27*).

#### Data inclusion criteria

For the polysomnographic records to be included into the data set for REM density measurements and SCI evaluation, individual sleep nights had to satisfy the following criteria: sleep efficiency of not less than 93%, with not less than 5 sleep cycles per night, and no signs of sleep apnea or other symptoms of sleep disorders (n=11 nights total). The last episode of the night was included in the analysis if REM sleep was not less than 25min long (n=5), to avoid episodes interrupted by wake onset.

#### NREMS and REMS episode assessment

NREMS-REMS cycles were defined by the succession of a NREMS episode of at least 10 min duration and a REMS episode of at least 3 min duration. No minimum criterion for REMS duration was applied for the completion of the last cycle. A NREMS episode was defined as the time interval between the first two epochs of stage 2 and the first occurrence of REMS within a cycle. A REMS episode was defined as the time interval between two consecutive NREMS episodes or as an interval between the last NREMS episode and the final awakening.

#### REMS intensity assessment

To quantify the number of eye movements during REM sleep we used the original methodology introduced by Aserinsky (10) to account for all the eye movements (REMs). Accordingly, the REMs were visually scored within 15-second intervals and the total number of eye movements per minute of REM sleep episode was calculated. All REMs detectable above the background noise were considered, irrespective of their amplitude, if they were present on both right and left electrooculography (EOG) channels simultaneously. All the stepwise saccades in the same direction of gaze were counted as separate eye movements.

#### The Invariant assessment

To assess the Sleep Cycle Invariant (SCI) for each sleep cycle, the REM density in each REMS episode was multiplied by the duration of prior NREMS sleep episode (i.e., within the same sleep cycle). To account for the first NREMS episode duration predicted to be, on average, curtailed by one quarter due to the position of sleep onset on the initial energy level of the Morse potential (*9*), the value was multiplied by 1.33 (4/3). The last sleep cycle was excluded from the analysis if the REMS episode duration was less than 25 min, suggesting it was interrupted by the morning awakening.

### (b) The “hybrid” analysis of SCI

*The NREMS episode duration* data were collected by Barbato & Wehr (*14*) in 11 healthy male volunteers, 20-34 years of age. The subjects were studied for 4 weeks, with regular activities over 10 hours of light and bedrest over 14 hours of darkness, when they were encouraged to sleep. The total of 308 sleep records were analyzed. The data used in the present study were obtained from Tables 2 in (*14*).

*The REMS intensity* (REM density) data were collected by Aserinsky (*11*) in 11 normal subjects, young males and females, identified as university students. Data for the initial night of a 54-h sleep-abundance protocol was used in the analysis (p. 550, in the text). The quantitative method of precise count of REMs was used to assess REM density, as originally developed by Aserinsky after he discover REM sleep.

### (c) The effect of the semi-quantitative method of REMS intensity evaluation on SCI

The NREMS episode duration and REMS episode intensity (evaluated using the semi-quantitative method), as reported by Barbato et al (*15*) (Table 2) were used in the analysis. In brief, the study was conducted in 8 healthy male volunteers (mean age = 29.0±4.5 years, range 23-34 years of age) over 208 sleep nights. The subjects were studied for 4 weeks, with regular activities over 10 hours of light and bedrest over 14 hours of darkness, when they were encouraged to sleep.

#### REMS intensity evaluation using the semi-quantitative Pittsburgh scale

The following detailed description of the semi-quantitative assessment of REMS intensity was provided by the authors (*15*): REM density was defined as total REM activity/REM duration. REM activity for each minute of REM was expressed on a 0-8 scale (mean of pairs of consecutive 30 sec REM epochs). According to this scale, 0 corresponded to no eye movements (EMs); 1, 1-2 EMs; 2, 3-5 EMs; 3, 6-9 EMs; 4, 10-14 EMs; 5, 15-20 EMs; 6, 21-26 EMs; 7, 27-32 EMs; and 8, 33 and over EMs.

#### NREMS episode duration assessment

In the original report (14), the NREM-REM cycles were analyzed and presented (Table 2) separately for two sub-groups. S-group 1: cycles not followed by period of wakefulness, NREM-REM-NREM (NR). S-group 2: cycles followed by wakefulness, NREM-REM-W (W). In Fig. 2d, we show the results for only Sub-group 1, i.e., for complete cycles only. When SCI was evaluated for the sub-group 2, the product of NREMS episode duration and REMS episode intensity was even further away from an invariant than in sub-group 1 (not shown).

### (d) Statistical assessment of SCI deviations from a constant value (Fig. 2B)

To test whether SCI deviates from a constant value across multiple sleep cycles, a linear mixed effect model was used. Specifically, we compared two models: H_1_ in which SCI value was modeled as a function of sleep cycle, and H_0_ in which SCI was modeled as a constant across all cycles. Both models included subject-specific intercept as a random effect, and treated cycle as a categorical variable. The models were fit on the 11 subjects and ANOVA test was used to test whether H_1_ explained significantly more deviance than H_0_. P value of 0.92 indicates that no significant deviation from a constant model was observed in the data. The calculations were carried out using lme4 package in R.

